# Population coding of reward prediction errors through opponent organization in the fronto parietal network

**DOI:** 10.1101/769869

**Authors:** Nicholas C. Foley, Michael Cohanpour, Mulugeta Semework, Sameer A. Sheth, Jacqueline Gottlieb

**Affiliations:** Department of Neuroscience, Columbia University; Zuckerman Mind Brain Behavior Institute, Columbia University; Department of Neurosurgery, Baylor College of Medicine; The Kavli Institute for Brain Science, Columbia University

## Abstract

Computing expectancy violations is essential for decision making and cognitive functions, but its neural mechanisms are incompletely understood. We describe a novel mechanism by which prefrontal and posterior parietal neurons encode reward prediction errors (RPEs) in their population but not single-neuron activity. Simultaneous recordings of neural populations showed that both areas co-activated information about experienced and expected rewards in a precise opponent organization. Neurons encoding expected rewards with positive (negative) scaling were reactivated simultaneously with those encoding experienced rewards with negative (positive) scaling. This opponent organization was mirrored in polarity-dependent noise correlations. Moreover, it extended to two types of expectancy information – based on task-relevant visual cues and statistically irrelevant reward history - allowing decoding of signed and unsigned RPE in two reference frames. Frontal and parietal areas implement canonical computations that facilitate contextual comparisons and the readout of multiple types of expectancy violations to flexibly serve behavioral goals.

## Introduction

Theoretical models of computational rationality postulate that utility-relevant outcomes, such as punishments or rewards, modulate not only our choices of action but also our allocation of cognitive resources^1, 2^. At both the neural and behavioral levels, cognitive resources – of attention, learning and memory – are allocated to events that have not been successfully predicted and violate expectations. Consistent with this view, the fronto-parietal executive network, which directs information processing in pursuit of behavioral goals, is recruited by response conflict, learning and exploration – i.e., tasks that have uncertainty and are likely to generate expectancy violations^3^.

Despite the importance of expectancy violations – or “prediction errors” – in regulating cognitive functions, their neural mechanisms are incompletely understood. By far the most intensively investigated responses are those of midbrain dopamine (DA) cells, which encode signed reward prediction error (RPEs) that closely resemble the canonical RPEs in artificial reinforcement learning algorithms. DA cells encode positive and negative RPE with significant excitation and inhibition for outcomes that are, respectively, better or worse than expected^4^. Circuit-level investigations suggest that these responses are computed by individual cells through neural subtraction of convergent excitatory and inhibitory inputs signaling, respectively, experienced and expected rewards^5^.

But, in contrast to the homogeneous responses in DA cells, neurons in other parts of the brain integrate responses to expected and experienced outcomes in a much more heterogeneous fashion and typically do not seem to encode RPEs^5^. Kennerley et al^6^ reported RPE-like responses in a minority of cells in the monkey dorsal anterior cingulate cortex (dACC), a node of the executive network that receives dopaminergic input and is involved in value-based regulation^7^. The RPE-encoding cells in the dACC showed anti-correlated value responses during expectancy and outcome epochs, consistent with a subtractive mechanism like that found in DA cells. However, the same authors reported that RPE signals are absent in the dorso-lateral prefrontal cortex (dlPFC), a lateral fronto-parietal area that is anatomically connected with the dACC despite the presence of robust reward-related activity^6, 8, 9^. Likewise, parietal areas connected with the dlPFC – specifically, area 7A on the inferior parietal gyral surface^10–13^ - have reward-related activity^14^, but it is unknown whether or how they encode RPEs.

Understanding whether and how structures beyond DA cells contribute to expectancy violations is important because, in natural tasks, multiple types of expectancy violations influence decisions and cognitive functions. Well-known distinctions are between model-based and model-free RPEs – calculated, respectively, based on task-relevant expectancies or irrelevant reward history^15, 16^, between surprises in appetitive outcomes or sensory observations^17^, and between surprises relative to specific objects or features^18–21^. Thus, it is important to understand how different types of expectancy violations are encoded in areas involved in decision making and cognitive functions.

To examine these questions, we simultaneously recorded from dlPFC and 7A cells while monkeys performed an instructed saccade task in which they received probabilistic rewards signaled by visual cues. We describe a novel mechanism by which dlPFC and 7A cells signal RPEs in the population but not single neuron activity. This population response was mediated by an opponent organization of cells encoding value with different polarity. Neurons that increased (decreased) firing in function expected rewards were co-activated with those that decreased (increased) firing as a function of the experienced rewards. Opponent organization encoded RPEs in two reference frames – based on task-relevant visual cues and statistically irrelevant reward history - suggesting that the fronto-parietal network may provide flexible readout of different types of expectancy violations for different behavioral goals.

## Results

### Task and behavior

Two monkeys performed a visually guided saccade task to obtain probabilistic rewards signaled by visual cues (**Fig. 1A**). Before neural recordings began, the monkeys were familiarized with 20 visual cues signaling 7 levels of EV through 10 distinct combinations of reward magnitude and probability (**Fig. 1B**). To perform the saccade task, the monkeys achieved central fixation, followed by the presentation of a randomly selected reward cue, a 600 ms memory period and the target for the instructed saccade (**Fig. 1A**). After making the required saccade, the monkeys received the outcome - reward omission or a reward with variable magnitude according to the contingencies signaled by the cue. A stereotyped tone marked the end of the post-saccadic hold period/onset of the outcome epoch on reward and no-reward trials. The cue and target locations were independently randomized, so that the cue was not predictive of the optimal action, and cues were equated for discriminability and counterbalanced across monkeys, with 2 cues assigned to each reward contingency (**Fig. S1A**). These features of the task design allowed us to examine reward-related responses independently of visual confounds, learning or the planning of instrumental actions.

**Figure 1.**
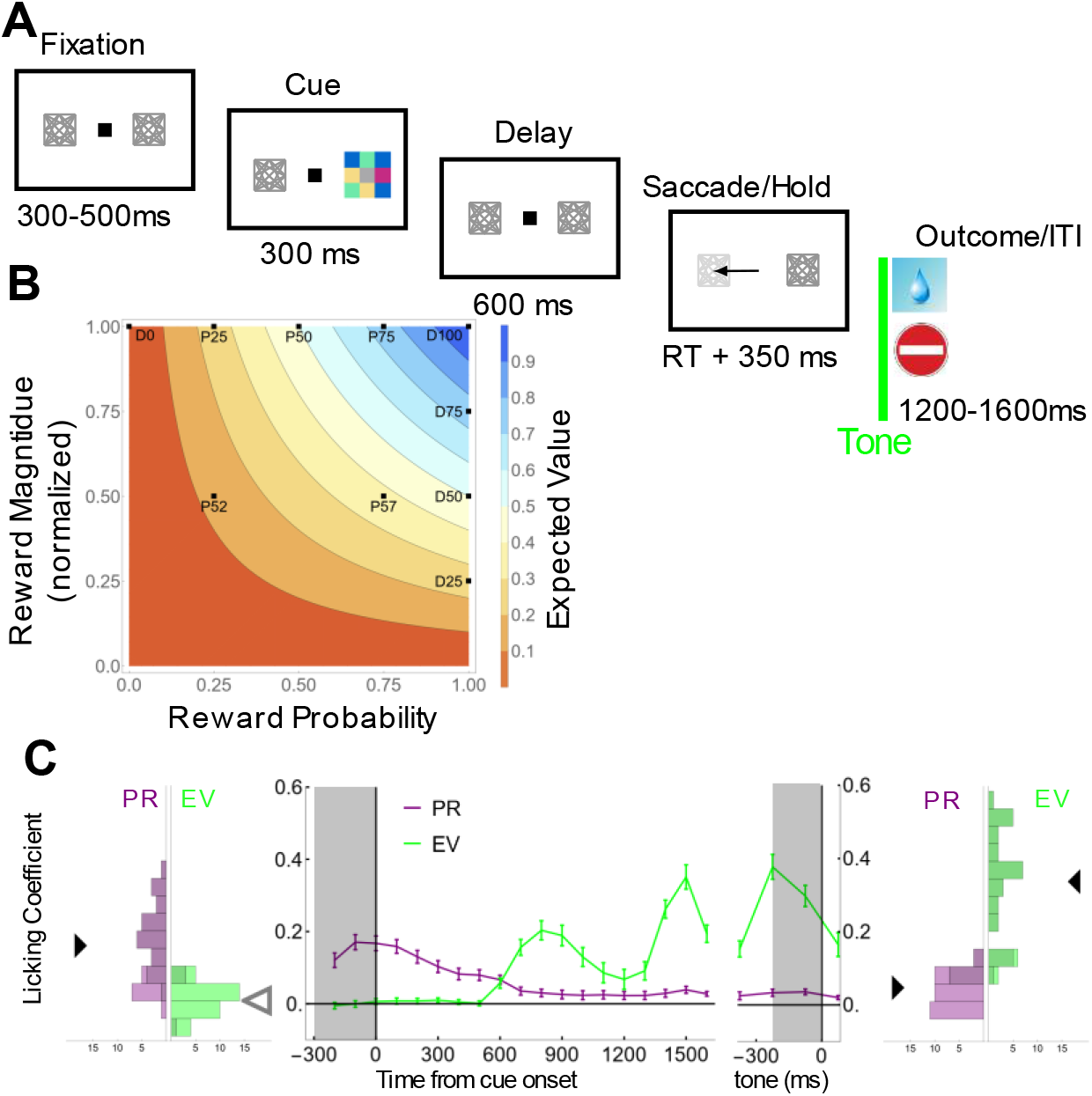
**(A) Task** stages. Monkeys fixated a central point (small black square) flanked by two placeholders (gray squares) and were shown a cue at a randomly selected placeholder location. After a delay period, a placeholder brightened, revealing the target for the required saccade. After the required saccade and a hold period, the monkeys received the outcome (reward (drop) or no reward; stop cartoon). Outcome period onset was marked by a tone and was followed by an intertrial interval (ITI). **(B) The set of reward contingencies** comprised 10 unique combinations of reward probability and magnitude (black points) with 7 levels of EV (colors indicate areas of constant EV). “P” cues indicated rewards with maximal size at 0.25, 0.5 or 0.75 probability and rewards with half size at 0.25 and 0.75 probability (P52 and P57, respectively. “D” cues indicated deterministic rewards of different magnitudes delivered deterministically with 0 or 100% probability. **(C) Licking is sensitive to EV and PR.** Marginal histograms show the distributions of regression coefficients estimating the effects of PR (purple) and EV (green) across sessions in the 200 ms windows before cue (left) and tone onset (right). Triangles show the means of the distributions, with filled colors indicating p < 0.05 relative to 0. The center panel shows the time-resolved coefficients (mean and SEM across sessions in 100 ms consecutive bins).

The monkeys’ anticipatory licking increased monotonically with the cue-signaled EV, confirming that they were familiar with and cognizant of the predictions made by the cues **(Fig. 1C**, green). Strikingly, even though prior rewards were statistically irrelevant to the current trial’s outcome (cues were trialwise randomized), the monkeys were strongly sensitive to the magnitude of the reward on the previous trial (**Fig. 1C**, purple). The effects of prior rewards (PR) were significant in 79% of individual sessions during the fixation period preceding the cue (**Fig. 1C**, linear regression analysis (*Methods*, Eq. 2); average PR coefficients, monkey 1: 0.21±0.006, p < 10^−10^; monkey 2: 0.04±0.01, p = 0.0001) and remained significant throughout the expectancy period when they coexisted with the effects of EV (**Fig. 1C**, marginal histograms on the right; PR coefficients 200 ms before outcome: monkey 1: 0.027±0.007, p = 0.0005; monkey 2: 0.054±0.01, p = 10^−6^; EV coefficients: monkey 1: 0.21±0.01, p < 10^−10^ relative to 0; monkey 2: 0.05±0.015, p = 0.0006). PR sensitivity was driven mostly by the immediately preceding reward with no significant influence of earlier trials (*Methods, Eq. 4*; PR coefficient for trial −1, 0.75±0.02 for monkey 1, p < 10^−10^; 0.65±0.04 for monkey 2, p < 10^−5^; trials −2 to −5, all p > 0.2). We found no evidence that the monkeys were using PR information to update their expectations about probabilistic cues (**Fig. S1B**), suggesting that they used their previously learned representations of the cues. Thus, the marked effect of PR occurred by default, independently of value updating or its relevance to the current trial EV.

### Frontal and parietal neurons encode expected and experienced outcomes

We recorded single neuron activity using multi-electrode “Utah” arrays implanted in the pre-arcuate dlPFC and in the OPT portion of area 7A activity (n = 1,298 and 736 cells, respectively; **Fig. S2**). Neurons in both areas carried information about EV, PR and the current reward (CR, the outcome that the monkeys received). We describe these responses and provide detailed statistical summaries in **Tables 1** and **2**.

**Table 1.**
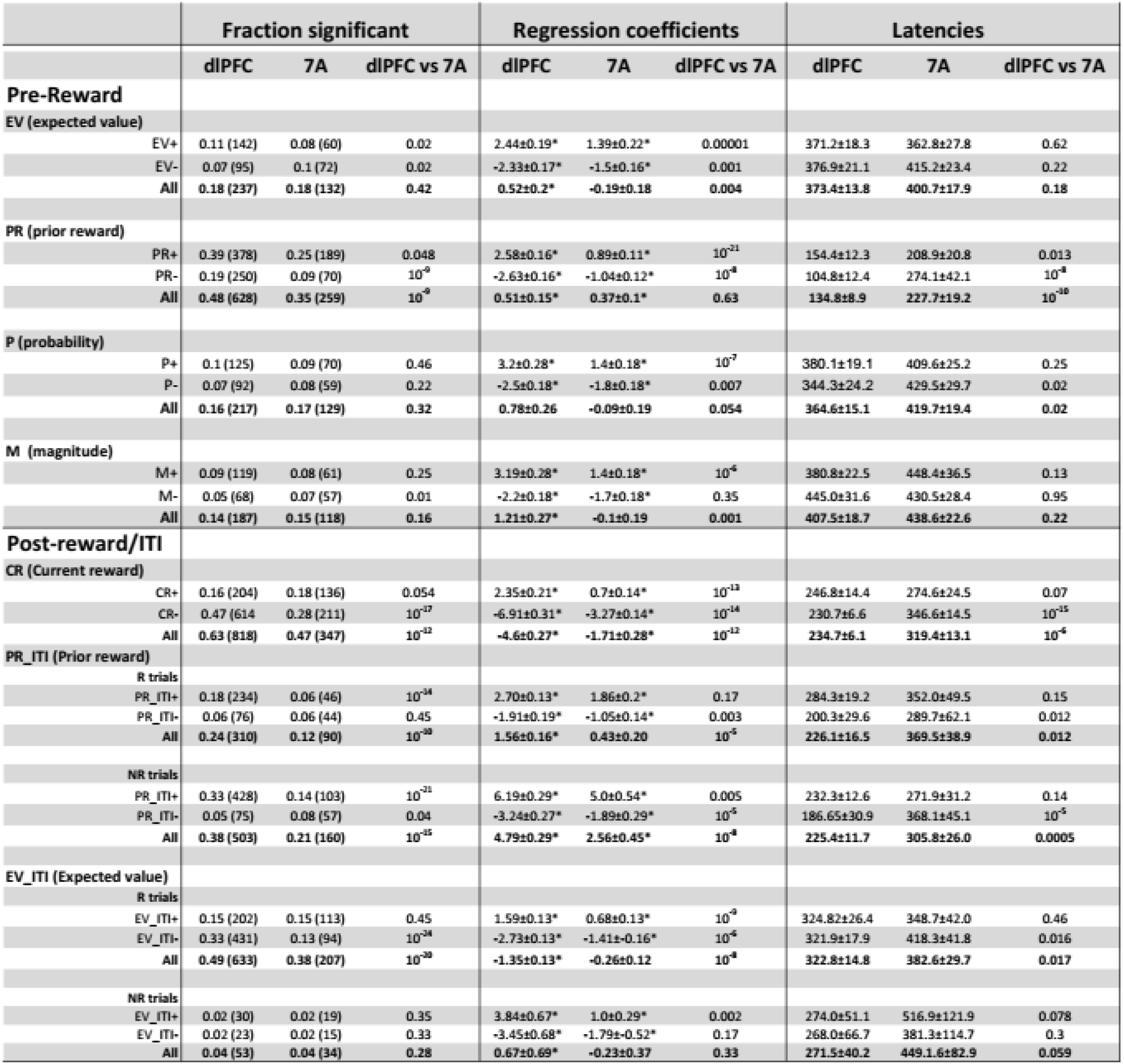
Neural modulations and area comparisons. The major vertical sections show, from left to right, the proportion of cells with significant effects (fraction (number)), the regression coefficient across the significant cells (mean±sem) and the latencies for significant cells (mean±sem). Each section gives the data for dlPFC and 7A, and the p-value for a statistical comparison across the two areas (left column, z-test of proportions; center and right column, two-sample tests as indicated in the text). The major horizontal sections show the signals discussed in the text, for positive encoding cells (+), negative encoding cells (−), and all the significant cells. EV, P and M effects are based on analyses in the 300 – 900 ms after cue onset; PR selectivity is based on 0 – 1,000 ms after fixation onset; and all the post-reward (ITI) effects were measured between 200 and 1,200 ms after tone onset. The analyses are based on the full data set (1,298 neurons in the dlPFC and 736 in 7A).

|  | Fraction significant |  |  | Regression coefficients |  |  | Latencies |  |  |
| --- | --- | --- | --- | --- | --- | --- | --- | --- | --- |
|  | dIPFC | 7A | dIPFC vs 7A | dIPFC | 7A | dIPFC vs 7A | dIPFC | 7A | dIPFC vs 7A |
| **Pre-Reward** |  |  |  |  |  |  |  |  |  |
| **EV (expected value)** |  |  |  |  |  |  |  |  |  |
| EV+ | 0.11 (142) | 0.08 (60) | 0.02 | 2.44±0.19* | 1.39±0.22* | 0.00001 | 371.2±18.3 | 362.8±27.8 | 0.62 |
| EV- | 0.07 (95) | 0.1 (72) | 0.02 | -2.33±0.17* | -1.5±0.16* | 0.001 | 376.9±21.1 | 415.2±23.4 | 0.22 |
| All | 0.18 (237) | 0.18 (132) | 0.42 | 0.52±0.2* | -0.19±0.18 | 0.004 | 373.4±13.8 | 400.7±17.9 | 0.18 |
| **PR (prior reward)** |  |  |  |  |  |  |  |  |  |
| PR+ | 0.39 (378) | 0.25 (189) | 0.048 | 2.58±0.16* | 0.89±0.11* | 10<sup>-21</sup> | 154.4±12.3 | 208.9±20.8 | 0.013 |
| PR- | 0.19 (250) | 0.09 (70) | 10<sup>-9</sup> | -2.63±0.16* | -1.04±0.12* | 10<sup>-8</sup> | 104.8±12.4 | 274.1±42.1 | 10<sup>-8</sup> |
| All | 0.48 (628) | 0.35 (259) | 10<sup>-9</sup> | 0.51±0.15* | 0.37±0.1* | 0.63 | 134.8±8.9 | 227.7±19.2 | 10<sup>-18</sup> |
| **P (probability)** |  |  |  |  |  |  |  |  |  |
| P+ | 0.1 (125) | 0.09 (70) | 0.46 | 3.2±0.28* | 1.4±0.18* | 10<sup>-7</sup> | 380.1±19.1 | 409.6±25.2 | 0.25 |
| P- | 0.07 (92) | 0.08 (59) | 0.22 | -2.5±0.18* | -1.8±0.18* | 0.007 | 344.3±24.2 | 429.5±29.7 | 0.02 |
| All | 0.16 (217) | 0.17 (129) | 0.32 | 0.78±0.26 | -0.09±0.19 | 0.054 | 364.6±15.1 | 419.7±19.4 | 0.02 |
| **M (magnitude)** |  |  |  |  |  |  |  |  |  |
| M+ | 0.09 (119) | 0.08 (61) | 0.25 | 3.19±0.28* | 1.4±0.18* | 10<sup>-6</sup> | 380.8±22.5 | 448.4±36.5 | 0.13 |
| M- | 0.05 (68) | 0.07 (57) | 0.01 | -2.2±0.18* | -1.7±0.18* | 0.35 | 445.0±31.6 | 430.5±28.4 | 0.95 |
| All | 0.14 (187) | 0.15 (118) | 0.16 | 1.21±0.27* | -0.1±0.19 | 0.001 | 407.5±18.7 | 438.6±22.6 | 0.22 |
| **Post-reward/ITI** |  |  |  |  |  |  |  |  |  |
| **CR (Current reward)** |  |  |  |  |  |  |  |  |  |
| CR+ | 0.16 (204) | 0.18 (136) | 0.054 | 2.35±0.21* | 0.7±0.14* | 10<sup>-13</sup> | 246.8±14.4 | 274.6±24.5 | 0.07 |
| CR- | 0.47 (614) | 0.28 (211) | 10<sup>-17</sup> | -6.91±0.31* | -3.27±0.14* | 10<sup>-14</sup> | 230.7±6.6 | 346.6±14.5 | 10<sup>-15</sup> |
| All | 0.63 (818) | 0.47 (347) | 10<sup>-12</sup> | -4.6±0.27* | -1.71±0.28* | 10<sup>-12</sup> | 234.7±6.1 | 319.4±13.1 | 10<sup>-6</sup> |
| **PR_ITI (Prior reward)** |  |  |  |  |  |  |  |  |  |
| **R trials** |  |  |  |  |  |  |  |  |  |
| PR_ITI+ | 0.18 (234) | 0.06 (46) | 10<sup>-34</sup> | 2.70±0.13* | 1.86±0.2* | 0.17 | 284.3±19.2 | 352.0±49.5 | 0.15 |
| PR_ITI- | 0.06 (76) | 0.06 (44) | 0.45 | -1.91±0.19* | -1.05±0.14* | 0.003 | 200.3±29.6 | 289.7±62.1 | 0.012 |
| All | 0.24 (310) | 0.12 (90) | 10<sup>-30</sup> | 1.56±0.16* | 0.43±0.20 | 10<sup>-5</sup> | 226.1±16.5 | 369.5±38.9 | 0.012 |
| **NR trials** |  |  |  |  |  |  |  |  |  |
| PR_ITI+ | 0.33 (428) | 0.14 (103) | 10<sup>-21</sup> | 6.19±0.29* | 5.0±0.54* | 0.005 | 232.3±12.6 | 271.9±31.2 | 0.14 |
| PR_ITI- | 0.05 (75) | 0.08 (57) | 0.04 | -3.24±0.27* | -1.89±0.29* | 10<sup>-5</sup> | 186.65±30.9 | 368.1±45.1 | 10<sup>-5</sup> |
| All | 0.38 (503) | 0.21 (160) | 10<sup>-15</sup> | 4.79±0.29* | 2.56±0.45* | 10<sup>-8</sup> | 225.4±11.7 | 305.8±26.0 | 0.0005 |
| **EV_ITI (Expected value)** |  |  |  |  |  |  |  |  |  |
| **R trials** |  |  |  |  |  |  |  |  |  |
| EV_ITI+ | 0.15 (202) | 0.15 (113) | 0.45 | 1.59±0.13* | 0.68±0.13* | 10<sup>-9</sup> | 324.82±26.4 | 348.7±42.0 | 0.46 |
| EV_ITI- | 0.33 (431) | 0.13 (94) | 10<sup>-34</sup> | -2.73±0.13* | -1.41±0.16* | 10<sup>-6</sup> | 321.9±17.9 | 418.3±41.8 | 0.016 |
| All | 0.49 (633) | 0.38 (207) | 10<sup>-30</sup> | -1.35±0.13* | -0.26±0.12 | 10<sup>-8</sup> | 322.8±14.8 | 382.6±29.7 | 0.017 |
| **NR trials** |  |  |  |  |  |  |  |  |  |
| EV_ITI+ | 0.02 (30) | 0.02 (19) | 0.35 | 3.84±0.67* | 1.0±0.29* | 0.002 | 274.0±51.1 | 516.9±121.9 | 0.078 |
| EV_ITI- | 0.02 (23) | 0.02 (15) | 0.33 | -3.45±0.68* | -1.79±0.52* | 0.17 | 268.0±66.7 | 381.3±114.7 | 0.3 |
| All | 0.04 (53) | 0.04 (34) | 0.28 | 0.67±0.69* | -0.23±0.37 | 0.33 | 271.5±40.2 | 449.1±82.9 | 0.059 |

**Table 2.** Correlations between regression variables. Each entry shows the Spearman correlation between the regression coefficients of a pair of variables (measured as in Table 1) across all the neurons in each area. White/gray shading indicates signals estimated during, respectively, the pre-reward and post-reward epochs. * p < 0.05; **p < 0.01; **\*\*\*p < 0.0001**

| dlPFC |  |  |  |  |  |  | 7A |  |  |  |  |  |
| --- | --- | --- | --- | --- | --- | --- | --- | --- | --- | --- | --- | --- |
|  | EV | P | M | CR | PR_ITI | EV_ITI | EV | P | M | CR | PR_ITI | EV_ITI |
| PR | 0.06* | 0.06* | 0.04 | -0.066* | 0.1** | 0.03 | -0.05 | -0.08* | 0.02 | 0.007 | 0.25*** | 0.05 |
| EV |  | 0.5*** | 0.69*** | -0.078* | 0.08* | -0.01 |  | 0.47*** | 0.68*** | -0.06 | -0.03 | 0.04 |
| P |  |  | 0.85*** | 0.02 | 0.11* | -0.02 |  |  | 0.45*** | -0.01 | -0.007 | -0.06 |
| M |  |  |  | 0.037 | 0.08* | -0.0003 |  |  |  | 0.046 | -0.02 | 0.05 |
| CR |  |  |  |  | -0.04 | -0.01 |  |  |  |  | -0.09* | 0.11* |
| PR_ITI |  |  |  |  |  | -0.39*** |  |  |  |  |  | -0.13* |

In the cue and delay epochs, approximately 18% of cells in both areas encoded the trial’s EV **Fig. 2A**; **Table 1**). EV selectivity was above and beyond any selectivity to the appearance and location of the visual cue (regression analysis, *Methods*, eq. 5) and included cells with positive (EV+) and negative scaling (EV−) – i.e., increases or decreases of firing at higher EV (**Fig. 2A**; **Table 1**). EV sensitive cells were equally prevalent in both areas but, among the sensitive cells, the dlPFC had higher coefficients and a higher proportion of EV+ cells relative to 7A (**Fig. 2A**; **Table 1**).

**Figure 2.**
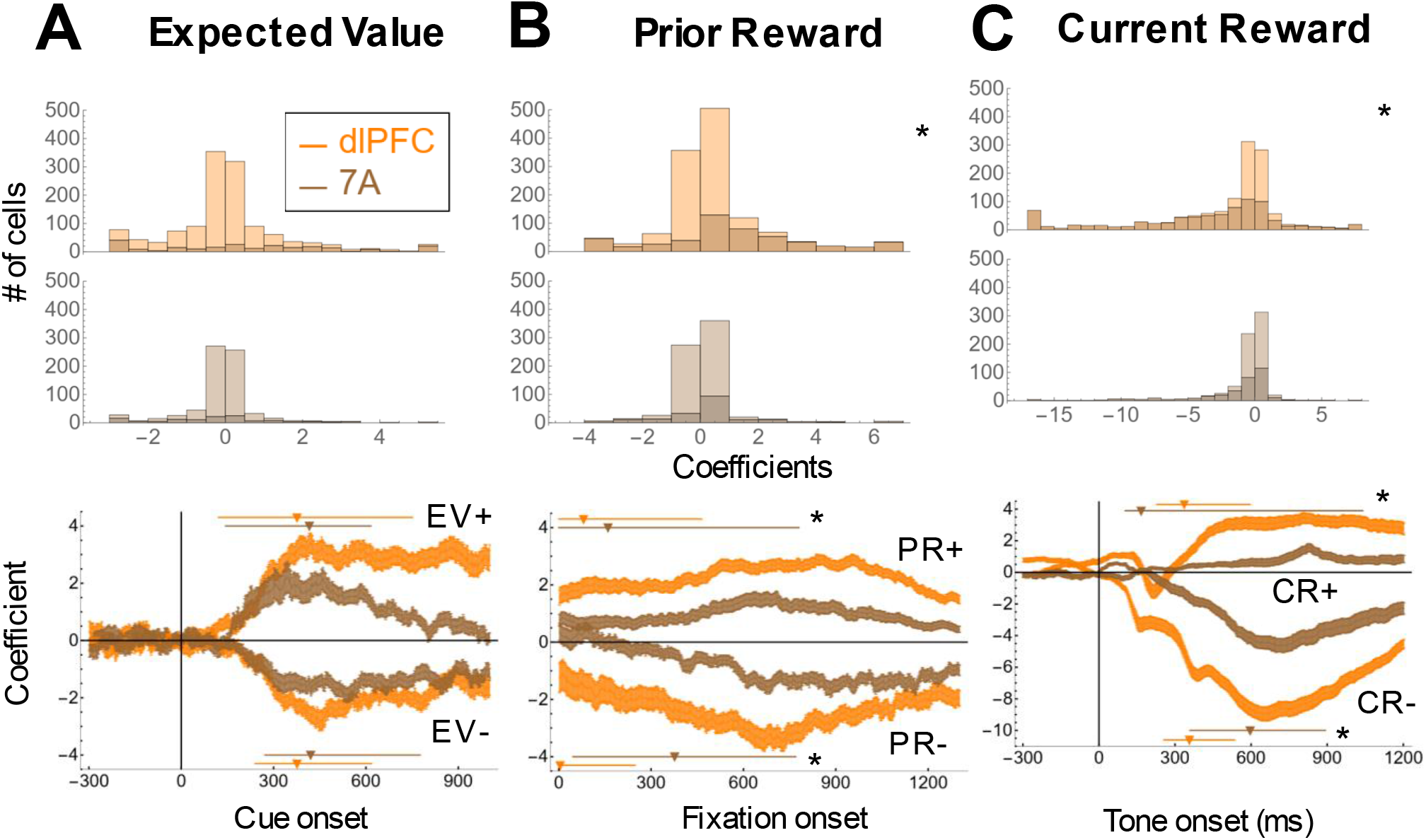
dlPFC and 7A neurons are sensitive to EV, PR and CR. **(A) EV selectivity.** Top: Histograms of EV coefficients during the cue and delay periods. Darker shading indicates cells with significant selectivity. Bottom: EV coefficients for cells with significant positive and negative scaling in a sliding window (mean and SEM, 50 ms bins and 2 ms steps aligned on cue onset). Triangles and lines show the median latencies and interquartile-intervals for the corresponding group. Stars indicate p < 0.05 for the latencies in dlPFC vs 7A. **(B) PR selectivity.** Analyses are aligned on fixation onset and shown in the same format as in A. **(C) CR selectivity** aligned on the onset of the outcome period (tone) in the same format as in A. The stars in **(B)** and **(C)** indicate p < 0.05 between dlPFC and 7A distributions.

In both areas, EV sensitivity arose at median latencies longer than 300 ms and was sustained throughout the delay period, confirming that it was not a mere visual response to the cues (**Fig. 2A**, **Table 1**). Moreover, EV coefficients were strongly correlated with the monkeys’ licking ehavior **Fig. S3A-C**) and with the neurons’ sensitivity to re ard pro a ility and magnitude (estimated on different subsets of trials; cf **Fig. 1B**; *Methods*, eq. 6,7; **Fig. S3D**). These strong correlations contrasted with the independence of EV coefficients from other reward modulations (**Table 2**), suggesting that the neurons integrated information about reward probability and magnitude to signal overall reward expectancy.

More than 33% of cells in each area encoded the size of the prior reward (PR; *Methods*, eq. 5; **Fig. 2B**), in a manner that was uncorrelated with sensitivity to EV (**Table 2**). PR sensitivity also showed positive or negative scaling (PR+ and PR−) – i.e., increases and decreases as a function of the prior trial reward – and was stronger and appeared earlier in the dlPFC relative to 7A (**Fig. 2B**, **Table 1**).

Several considerations showed that PR sensitivity was not merely a continuation of responses to the previous outcome. First, PR sensitivity arose with latencies of more than 100 ms after monkeys acquired fixation (earlier in dlPFC relative to 7A; **Fig. 2B**, bottom; **Table 1**). Second, classifiers trained to decode CR from the population activity in the outcome epoch performed poorly in decoding the same reward during the fixation period of the following trial, and vice versa, showing that the population representation o changed at a trial’s onset (**Fig. S4**). Finally, the representation of PR in the population response strongly resem led the representation in the monkeys’ licking ehavior **Fig. S3C**). Thus, the PR-related activity was an active recall of the recent reward history that was combined with EV to contri ute to the monkeys’ reward expectations.

The final and strongest response in both areas was evoked by the current reward (CR) – the size of the reward that the monkeys experienced (**Fig. 2C**). CR sensitivity was significant in 63% of dlPFC cells and 47% of 7A cells (*Methods*, eq. 8). CR responses also showed positive and negative scaling (CR+ and CR−), but were strongly biased toward negative scaling, with more than 60% of the CR sensitive cells in each area responding preferentially to *omission* rather than receipt of reward (CR = 0; **Fig. 2C**, **Table 1**). Although very robust in both areas, CR sensitivity was stronger in dlPFC, as indicated by the higher prevalence of significant selectivity (p < 10^−12^; **Fig. 2C**, **Table 1**) and, in the sensitive cells, stronger modulations and shorter latencies in the dlPFC relative to 7A (**Fig. 2C**, **Table 1**).

### Information about expected rewards is reactivated after the outcome

dlPFC neurons have been shown to recall information about visual cues at the time of outcome delivery, suggesting that they contribute to credit assignment mechanisms^22^. We found that the neurons showed an analogous recall of information in our task which, rather than signaling cue identity, conveyed information about PR and EV. These post-outcome representations of PR and EV (which we term PR_ITI and EV_ITI; *Methods*, eq. 8; **Fig. 3** and **Table 1**) were not mere continuations of delay-period activity, as coefficients for EV_ITI and PR_ITI were uncorrelated with those for, respectively, EV and PR (**Table 2**). Moreover, the accuracy of classifiers trained and tested in different time epochs declined at boundary set by outcome delivery (**Fig. 3**). Classifiers trained to decode PR or EV after outcome delivery were significantly less accurate at decoding the same variable in an epoch preceding the outcome, relative to an equally distant time window after the outcome (**Fig. 3**, right panels), showing that the population representation of PR and EV changed at the time of outcome delivery.

**Figure 3.**
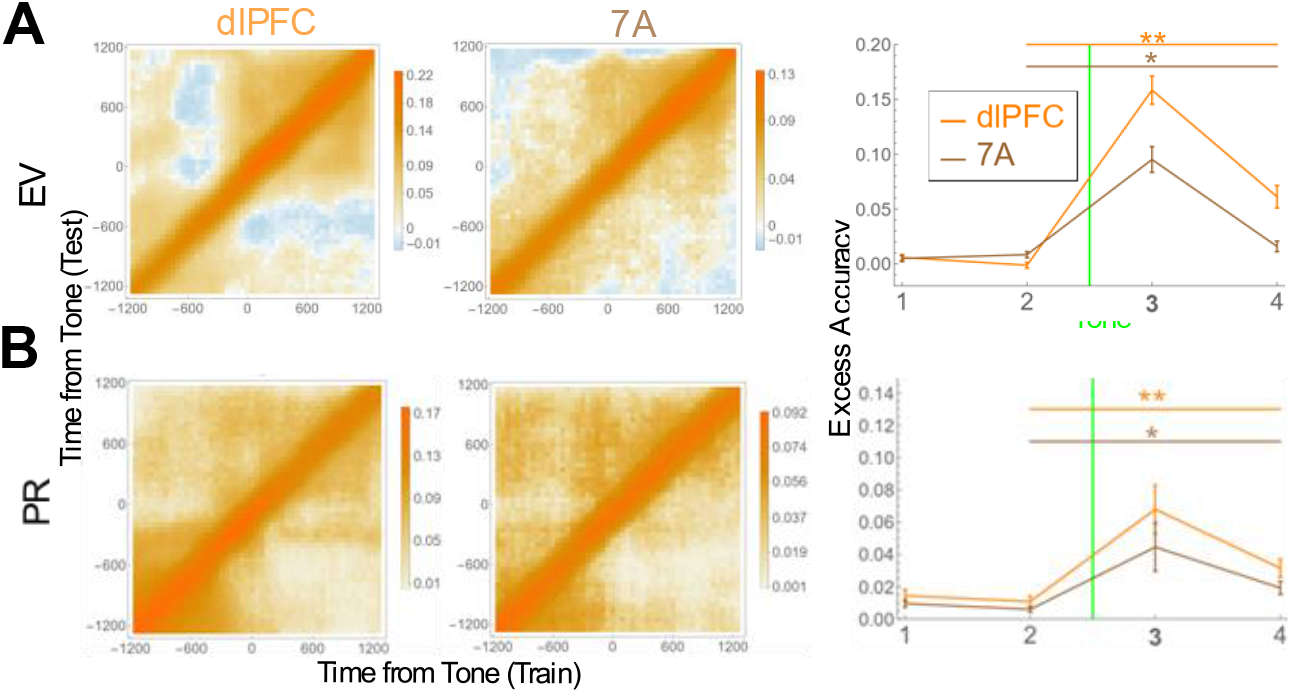
EV and PR information after the outcome has a different representation. **(A) Accuracy of logistic classification** trained and tested at different times before and after the outcomes. Logistic classifiers were trained on 300ms epochs, sliding at 50ms to decode EV from the neural populations, and tested on their accuracy in all other epochs In the left panels, the colors denote excess accuracy, decoding accuracy minus the accuracy from control data with randomly shuffled labels. The right panel shows classifiers trained at 0 to 600ms from tone and tested against neural data at −1200 to −600 (epoch 1), −600 to 0 (epoch 2), the training epoch, and 600 to 1200 (epoch 4) from the time of the tone (* p < 0.05; ** p < 0.001). **(B)** Decoding of PR, same conventions as in A

### Cells encoding experienced and expected outcomes show opponent organization

The reemergence of EV and PR sensitivity suggests that dlPFC and 7A neurons may integrate information about expected and experienced outcomes to encode RPE. The distinguishing signature of RPE encoding in individual cells is an anticorrelation between responses to expected and experienced outcomes^5, 6^. This negative correlation, however, was not present in our data, as there were no convincing negative correlations between coefficients of CR and EV_ITI, or CR an PR_ITI (**Table 2**).

At the population level, however, the newly emerged signals of PR_ITI and EV_ITI showed a precise opponent organization ith respect to the trial’s re ard and selectivity that resulted in an emphasis of less expected outcomes. Sensitivity for PR_ITI was predominantly *positive* (higher responses for larger prior rewards) and was stronger after *omission* relative to receipt of reward (**Fig. 4A**). In contrast, sensitivity for EV_ITI was predominantly *negative* and was stronger after reward *receipt* relative to omission (**Fig. 4B**). For PR_ITI, the pattern was very clear both in the subset of significant cells (**Table 1**) and in the entire population in both areas (dlPFC, PR_ITI coefficients after omission vs receipt: 4.79±0.29 vs 1.56±0.16; p < 10^−14^; 7A: 2.56±0.45 vs 0.43±0.2; p < 10^−11^). For EV_ITI, the dependence on outcome was not seen in 7A (EV_ITI coefficients of −0.26±0.12 vs −0.23±0.37, all neurons, p = 0.74) but was highly robust in the dlPFC (EV_ITI coefficients after reward receipt versus omission, −1.35±0.13 vs 0.67±0.69, all neurons, p < 10^−19^). Thus, an opponent organization resulted in emphasis on different unexpected outcomes in different reference frames - reward omissions that followed a large prior reward, and reward receipt that followed a lower cue-predicted EV.

**Figure 4.**
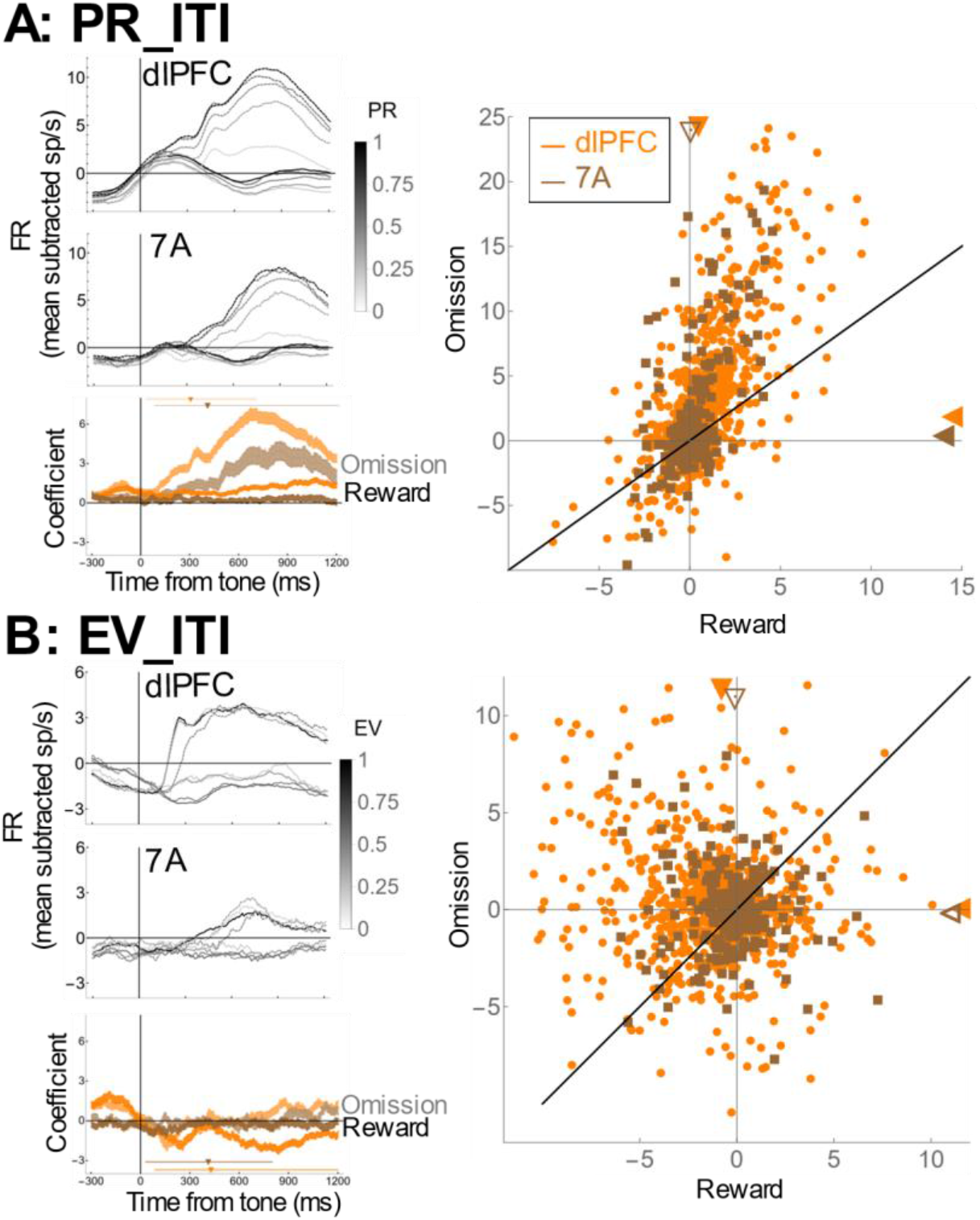
Opponent organization of expected and experienced outcome in firing rates. **(A) Across the population, the sensitivity to PR_ITI is positive and stronger after reward omission.** The top two left panels show average PSTHs for cells with PR_ITI sensitivity, aligned on tone onset, separated according to the outcome (reward receipt or omission) and PR (grayscale). The higher firing rates are trials in which rewards were omitted. For each outcome, the neurons have the strongest responses for the highest PR, and this effect is stronger after reward omission. The bottom left panel shows the PR_ITI coefficients from eq. 8 (sliding window of 50 ms width and 2ms steps). The traces show the mean and SEM for the sensitive cells in dlPFC (orange) and 7A (brown), separately for trials ending in reward (darker shade) and reward omission (lighter shading; orange: dlPFC; brown: 7A). The right panel is a comparison of regression coefficients for PR_ITI (*Methods*, eq. 8) on rewarded and unrewarded trials, for all the recorded cells. Each point is one cell. The closed triangles represent population mean (closed, p < 0.05 relative to 0, open, NS). **(B) Across the population, the sensitivity to EV_ITI is stronger on rewarded trials and predominantly negative.** Same conventions as in **A.**

We examined whether the opponent organization reflects patterns of functional connectivity by measuring noise correlations - the extent to which pairs of simultaneously recorded neurons have correlated fluctuations in their trial by trial responses^23^. We restricted this analysis to pre-cue activity and removing the influence of PR (*Methods*), ensuring that we estimate the neurons’ shared functional connectivity above and beyond that which generates their average task-related responses^23^. Consistent with the firing rate patterns, noise correlations were greatest among cells that encoded experienced and expected outcomes with opposing polarity (**Fig. 5**).

**Figure 5.**
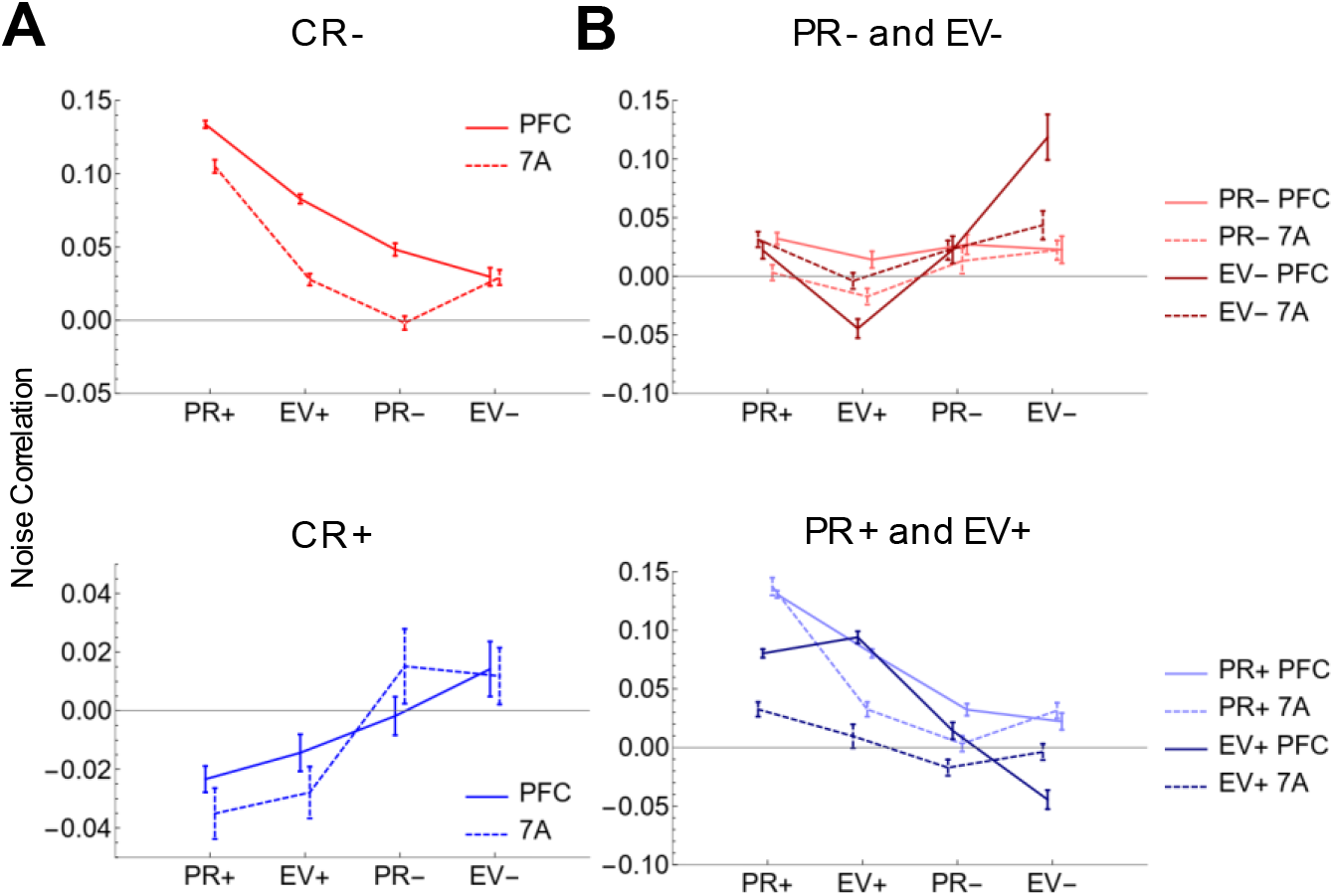
Opponent encoding in noise correlations. **(A)** Noise correlations computed in the 300ms prior to cue onset (PR removed) (significant cells from Figure 2 with a minimum firing rate of 4hz). CR− cells, top panel in red (solid PFC, dashed 7A), CR+ cells bottom panel in blue. CR− cells show opposite polarity correlation with EV+ and PR+ cells, while CR+ cells show weaker correlations in general and negative correlations with EV+ and PR+ cells. **(B)** Top panel: PR− (light red, solid PFC, dashed 7A; again from Figure 2, minimum 4hz firing rate) and EV− cells (dark red, solid PFC, dashed 7A) show low noise correlations with the exception of a putative EV− network. PR+ (light blue, solid PFC, dashed 7A) and EV+ (dark blue, solid PFC, dashed 7A) cells show strong positive correlations with the same polarity both within and between modalities.

The opponent organization of noise correlations was most evident in CR− cells, which had stronger noise correlations with PR+ and EV+ relative to PR− and EV− cells (**Fig. 5A**, top; 1-way Mann-Whitney, main effect of polarity, dlPFC, p = 10^−23^, 7A, p = 10^−13^). Moreover, CR− cells had stronger correlations with PR+ relative to EV+ cells (**Fig. 5A**, top; 1-way Mann-Whitney, PR+ vs EV+ dlPFC, p = 10^−66^, 7A, p = 10^−53^) consistent with the higher effect of PR than EV on reward omission responses (cf **Fig. 4A**). CR+ cells followed this opponency principle, having higher (more positive) noise correlations with PR− and EV− cells relative to PR+ and EV+ cells (**Fig. 5A**, bottom; 1-way Mann-Whitney, main effect of polarity (+ vs-, PR and EV together) dlPFC, p = 0.026, 7A, p = 0.12), resulting in a significant interaction between CR+ and CR− connectivity patterns (Friedman test, p < 0.01 in each area).

The opponent pattern of CR-sensitive cells differed greatly from that found among neurons encoding PR and EV, for which noise correlations were higher for neurons with similar encoding polarity (**Fig. 5B**). This pattern was strongest for PR+ and EV+ cells, which had stronger correlations with other positive polarity cells (**Fig. 5B**, bottom; 2-way Friedman ANOVA, dlPFC: main effect of polarity, p = 10^−35^, main effect of variable (PR vs EV) p = 10^−77^; 7A: main effect of polarity, p = 10^−21^, main effect of variable (PR vs EV) p = 10^−28^) whereas PR− and EV− cells tended to have more positive correlations with other negative polarity cells (**Fig. 5B**, top; 2-way Friedman ANOVA, dlPFC: main effect of polarity p = 0.13, main effect of variable (PR vs EV) p = 0.003; 7A: main effect of polarity (+ vs-) p = 0.48, main effect of variable (PR vs EV); p = 0.003). Thus, the opponent organization in the neural responses may arise from opponent patterns of functional connectivity among cells encoding expected and experienced outcomes.

### RPEs are decoded from population activity but not individual cells

To measure the information conveyed by this opponent organization, we explicitly compared the encoding of RPEs in individual neurons and population activity. Analysis of individual cells revealed no evidence that they consistently encoded RPE. Plotting firing rates as a function of RPE in two references frames (PR_RPE, the difference between the magnitude of the current and prior reward - and EV_RPE the difference between the reward magnitude and the trial’s EV) revealed no cluster that consistently scaled as a function of RPE (**Fig. S5**). On average across cells, a weak linear trend for PR_RPE was found only in the dlPFC and only for negative RPE on rewarded trials (**Fig. 6A**), reflecting the responses of small clusters of individual cells (**Fig. S5**; dlPFC: cluster 3, 24% of the cells; 7A: cluster 5, 14% of the cells). A consistent relation with EV_RPE was even more elusive overall and in individual clusters (**Fig. 6B**; **S5**).

**FIGURE 6.**
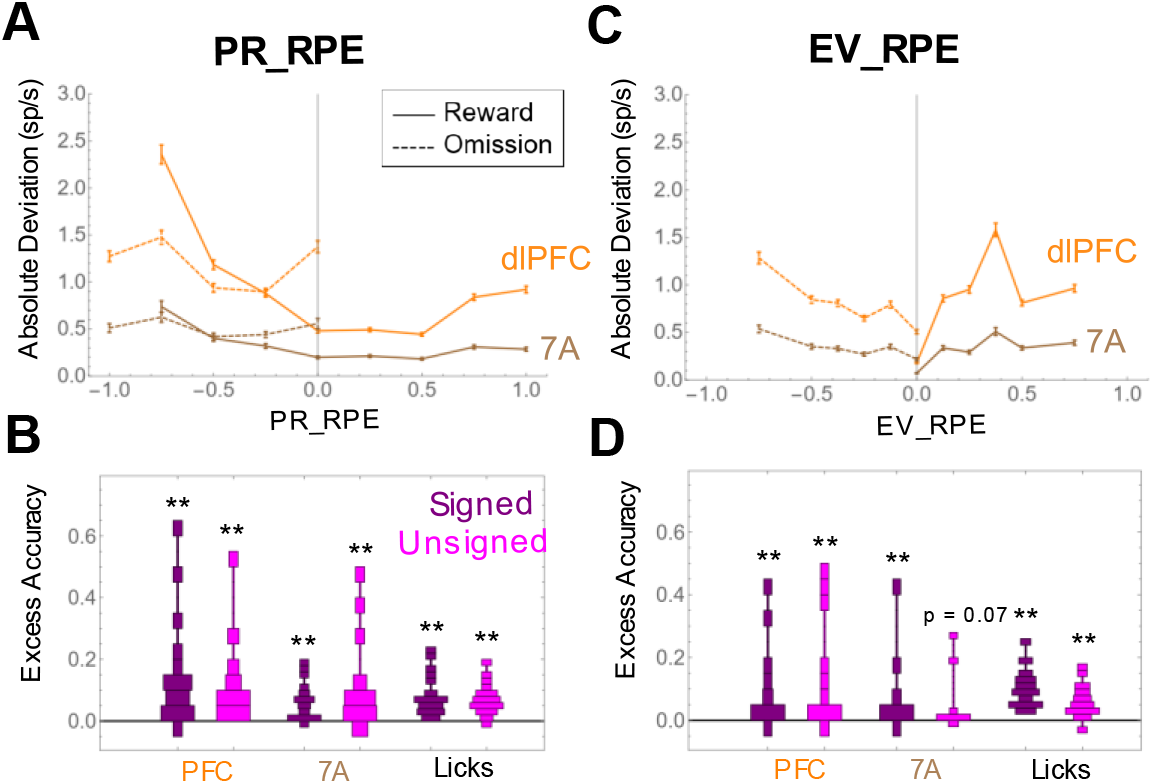
PR_RPE and EV_RPE are decoded from the population but not individual cells. **(A) Non-linear response to PR_RPE** The absolute deviation from the average firing rates as a function of PR_RPE, after reward receipt (solid) and omission (dashed). The points show the mean and SEM for all the cells in dlPFC (orange) and 7A (brown). **(B) Decoding of PR_RPE** Distributions of decoding accuracy (after subtraction of accuracy in shuffled trials) across the recording sessions, for dlPFC, 7A, and LR, and for signed and unsigned PR_RPE. (* p < 0.05; ** p < 0.001) **(C) Non-linear responses to EV_RPE** Same conventions as in **A**. **(D) Decoding of EV_RPE** Conventions as in **B**.

Despite the lack of encoding in individual cells, signed and unsigned PR_RPE and 7 EV_RPE could be decoded from the population response in both areas (**Fig. 6C, D**). A 3-way ANOVA of accuracy across sessions with factors of area (dlPFC vs 7A), reference frame (EV_RPE vs PR_RPE) and type of RPE (signed or unsigned) showed that decoding accuracy was above-chance in both areas but was higher in the dlPFC relative to 7A (p = 0.005; **Fig. 6C,D**). Decoding accuracy was similar for signed and unsigned RPE (p = 0.9). Remarkably, RPE was decoded more reliably if referenced to the statistically irrelevant reward history than to the task-relevant cue (p = 0.0003). Finally, PR_RPE and EV_RPE could e decoded rom the monkeys’ licking responses in both their signed and unsigned forms, showing that these quantities were behaviorally significant.

## Discussion

Neurons encoding expected and experienced rewards with positive and negative polarity have been reported in frontal cortical areas – including the dlPFC, dACC and orbitofrontal cortex^6, 8, 9^ – but their functional significance is not understood. Here we show that such neurons are also found in the inferior parietal area 7A that is anatomically connected with the dlPFC. Importantly, in both the frontal and parietal areas they are instrumental in constructing population-level representations of RPEs.

The representations of RPE we describe differ fundamentally from the well-known responses to RPE conveyed by midbrain DA cells. Individual DA cells, like a subset of dACC cells, explicitly encode RPE by virtue of convergent (anticorrelated) inputs signaling the values of expected and experienced outcomes^5, 6^. In contrast, responses to expected and experienced outcomes were uncorrelated in 7A and the dlPFC, and individual neurons in these areas did not scale linearly with RPE. Instead, both areas signaled RPEs in their population activity through the selective co-activation of classes of neurons with opposite polarity.

Large fractions of neurons in each area encoded the value of rewards that the monkeys received with *negative* polarity and were co-activated with neurons encoding the reward expected based on recent history with *positive* polarity. A smaller subset of neurons encoded the value of the rewards that the monkeys received with *positive* polarity and were co-activated with neurons encoding the reward expected based on the visual cue with *negative* polarity. Thus, a precise population-level opponent organization conveyed RPEs in two reference frames - relative to the task-relevant cues and the statistically irrelevant reward history. In both cases, signals of reward expectancy after the outcome were not merely passive continuations of those preceding the outcome but were actively recalled in distinct populations of cells in a pattern that was mirrored by polarity-dependent noise correlations. Together, the findings suggest that frontal and parietal areas implement several mechanisms that allow them to contextualize reward outcomes and detect those that violate expectations – including responses with different polarity, opponent specific functional connectivity, sustained short-term memories and the active recall of recent information.

The robust effects of reward history we observed were striking given that prior rewards were statistically irrelevant for the current trial expectations. Responses to recent rewards have been often reported in frontal and parietal areas and related to trial and error learning^22, 24, 25^, decision making on longer time scales^26, 27^ and adaptation to errors and conflict^28^. A few studies have noted that reward-history responses remain prominent even when conveying seemingly irrelevant information, as was the case in our task, but have not offered an interpretation of their functional significance^29, 30^.

Our results suggest two potentially related functions for default encoding of reward history. First, PR-related responses were correlated with the PR-induced bias in the monkeys’ reward expectations. This suggests they may contribute to the “hot-hand” allacy, a ell-known economic bias whereby decision makers expect that positive or negative outcomes happen in streaks independently of their true autocorrelations^31^. Second, the opponent organization of PR_ITI versus CR-related responses highlighted differences between successive outcomes – i.e., the temporal derivative of reward history. Thus, a key contribution of PR-related activity may be to the contextual monitoring of overall reward rates, which is essential for hierarchical decisions on longer time-scales (e.g., determining whether to persist with a task or whether (and when) to abandon the task)^32^. Together, these findings suggest that fronto-parietal responses to reward history may link two apparently disparate computations - value monitoring on longer time scales and spurious serial biases in reward expectations.

It was also remarkable that the decoding of RPEs referenced to reward history was stronger than that of RPEs referenced to the relevant visual cues. This counterintuitive observation is likely explained by the fact that monkeys were familiar with the probabilities signaled by the cues. The study by Asaad and colleagues^22^ showed that dlPFC neurons recall a representation of visual cues at the time of reward delivery, but only while monkeys learn to assign credit to one of several cues(ref). The reactivated responses we found encoded the value (not identity) of the visual cues, and may have contributed to updating the monkeys’ elie s a out the cue-signaled reward contingencies^33^. However, our highly trained monkeys relied on previously-learned associations without continuously updating cue probabilities, potentially explaining the weaker re-emergent EV-related activity.

Canonical RPE responses, such as those found in DA cells, are thought to unction as “critics” that provide plasticity signal for reinforcing connections associated with increases in reward expectations^4^. The fronto-parietal responses we found are distributed and implicit in population activity and do not seem well suited for a similar role. Instead, we propose that these responses point to more general fronto-parietal mechanisms of contextualizing ongoing events. By virtue of their functional properties, including long time-constants^34^ that allow sustained memories, and a functional connectivity structure that recruits reward-sensitive neurons in an opponent organization, fronto-parietal neural ensembles can implicitly and simultaneously encode multiple types of expectancy violations. Thus, these areas serve as a reservoir of information from which downstream areas can monitor expectancy violations in multiple reference frames to flexibly serve behavioral goals.

## Methods

### General methods

Data were collected from two adult male rhesus monkeys (Macaca mulatta; 9-12kg) using standard behavioral and neurophysiological techniques as described previously^35^. All methods were approved by the Animal Care and Use Committees of Columbia University and New York State Psychiatric Institute as complying with the guidelines within the Public Health Service Guide for the Care and Use of Laboratory Animals. Visual stimuli were presented on a MS3400V XGA high definition monitor (CTX International, INC., City of Industry, CA; 62.5by 46.5 cm viewing area). Eye position was recorded using an eye tracking system (Arrington Research, Scottsdale, AZ). Licking was recorded at 1 kHz using an in-house device that transmitted a laser beam between the tip of the juice tube and the monkey’s snout and generated a 5V pulse upon detecting interruptions of the beam when the monkey extended his tongue to obtain water.

### Task

A trial started with the presentation of two square placeholders (1° width) located along the horizontal meridian at 8° eccentricity to the right and left of a central fixation point (white square, 0.2° diameter). After the monkey looked maintained gaze on the fixation point for 300-500 ms (fixation window, 1.5-2° square) a randomly selected placeholders was replaced for 300 ms by a reward cue – a checkerboard pattern indicating the trial’s re ard contingencies (see **Fig. S1A** for detailed description of the visual appearance of the cues). After a 600ms delay period, the fixation point disappeared simultaneously with an increase in luminance of one of the placeholders (the target), whose location was randomized independently from that of the cue. If the monkey made a saccade to the target with a reaction time (RT) of 100 ms – 700 ms and maintained fixation within a 2.0-3.5° window for 350 ms, he received a reward with the magnitude and probability that had been indicated by the cue. An auditory tone (200 ms, 500 Hz) signaled the end of the post-saccadic hold period on all trials, providing a temporal marker for the onset of the outcome/ITI period whether a reward was received or omitted. Rewards, when delivered, were linearly scaled between 0.28 to 1.12mL. The ITI – from tone onset to the onset of the fixation point on the following trial lasted for 1200-1600 ms. Error trials (resulting from fixation breaks, premature, late or wrong-direction saccades) were immediately repeated until correctly completed, precluding the monkeys from aborting trials in which they anticipated lower rewards. Monkeys were extensively familiarized with the task and all the cues before recordings began.

### Neural recordings

After completing behavioral training, each monkey was implanted with two 48-electrode Utah arrays (electrode length 1.5 mm) arranged in rectangular grids (1 mm spacing; monkey 1, 7×7 mm, monkey 2, 5×10 mm) and positioned in the pre-arcuate portion of the dlPFC and the posterior portion of area 7A (**Figure S2**). Data were recorded using the Cereplex System (Blackrock, Salt Lake City, Utah) over 22 sessions spanning 4 months after array implantation in monkey 1, and 12 sessions spanning 2 months after implantation in monkey 2.

### Data analysis

Error trials were discarded and not considered further (13.7% in monkey 1, 14.3% in monkey 2). All statistical analyses were preceded by tests of normality and symmetry (p < 0.05). For univariate comparisons we used the Wilcoxon-signed-rank test if the symmetry criterion was met, and the Mann-Whitney U-test otherwise. Correlation coefficients were computed using the Spearman Rank test.

### Behavior

Eye position was digitized at 220 Hz, and saccade RT was defined using velocity and acceleration criteria^36^. While RT showed some effects of reward contingencies and spatial congruence between cue and target locations, these effects were not consistent and are not reported here.

Trial-by-trial licking rates (LR) were defined as the proportion of time spent licking in a time window of interest. To estimate the effects of PR we focused the analysis on pairs of consecutive correct trials. We used a 3-step hierarchical analysis (eq. 1-3) to separately estimate the influence of PR and EV on LR. In the first step we partitioned out the effect of the prior trial’s outcome *type* (reward receipt of omission):

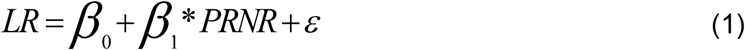

where RNR is an indicator equal to 1 if the prior trial was reward and 0 otherwise. In the second step used the residuals from eq. 1 to estimate the effect of PR:

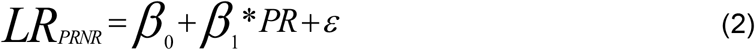

where *LR*_*PRNR*_ are the residuals from eq. 1, and PR is the magnitude of the prior reward (0, 0.25, 0.5, 0.75, 1). hus, the coe icients e report estimate the monkeys’ sensitivity to the size o the prior reward above and beyond the mere presence of a reward. Finally, in the third step we used the residuals from eq. 2 to estimate the effect of EV:

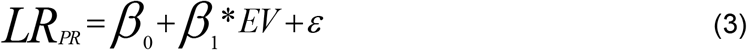

Where *LR*_*PR*_ are the residuals from equation 2, and EV is the expected value of the current cue (0, 0.125, 0.25, 0.375, 0.5, 0.75, 1).

To assess the temporal window over which reward history exerted effects, licking was regressed against the value of the prior reward (0, 0.25, 0.5, 0.75, 1) for the 5 previous trials, omitting error trials:

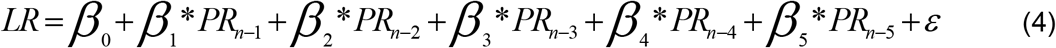

### Neural responses

Raw spikes were sorted offline using WaveSorter^37^ and analyzed with MatLab (MathWorks, Natick, MA) and Mathematica (Champaign, IL). Only neurons with waveforms clearly separated from noise were included in the analysis. All neural analyses were computed on unsmoothed firing rates (FR) that had been normalized within each cell by subtracting the FR averaged across the entire epoch from fixation onset until the end of the ITI.

We used regression analyses to measure the sensitivity to EV, PR and CR. All regressors ranged between 0 and 1, and took values of [0, 0.125, 0.25, 0.375, 0.5, 0.75, 1] for EV, and [0, 0.25, 0.5, 0.75, 1] for PR and CR. Coefficients are reported in units of sp s^−1^, and sensitive cells are defined as those showing a coefficient with p-value < 0.05.

We included cue location (CL) and target location (TL) as nuisance regressors (coded as 0 or 1 for the hemifield that was, respectively, ipsilateral or contralateral to the recording site), ensuring that we estimate sensitivity to reward variables independently of spatial coding or reward x space interactions.

To estimate the effects of PR and EV (**Fig. 3** and **Table 1**) we fit FR using the equation:

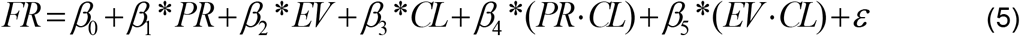

We defined a cell as being PR-sensitive if it showed a significant β_1_ coefficient in the interval 0 – 1,000 ms after fixation point onset, and EV-sensitive if it showed a significant β_2_ coefficient in the delay period (300 - 900 ms after cue onset).

To estimate the effects of reward probability (P) we fit firing rates on trials with probabilistic cues using the equation:

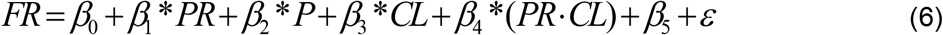

To estimate the effects of reward magnitude (RM) we fit firing rates on trials using deterministic cues using the equation:

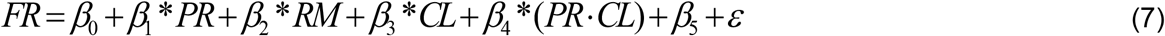

We defined a cell as being sensitive to probability or magnitude if it had a significant coefficient in the delay period (same interval as that used to measure sensitivity to EV).

To estimate the effects of CR, EV_ITI and PR_ITI we fit FR using the equation:

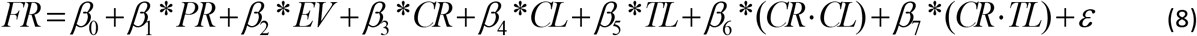

To estimate sensitivity to CR we applied this equation to all trials and defined a cell as being CR-sensitive if it showed a significant β_3_ coefficient in the interval 200 – 1,200 ms after tone onset.

To measure the sensitivity to EV_ITI and PR_ITI while accounting for the asymmetry of these modulations, we re-applied eq. 8 separately to trials ending in reward and reward omission. (For the latter trials, the CR term was dropped as it was always equal to 0). We defined a cell as being “sensitive” ased on the trials ith the strongest modulations i.e., i it had a significant β_2_ coefficient for EV_ITI on rewarded trials, or a significant β_1_ coefficient for PR_ITI on unrewarded trials).

To estimate effect latencies, we focused on the subset of cells that were sensitive for each factor and analyzed them with reduced models that included only an intercept term and the regressor of interest and was applied in a 50 ms window stepped by 2 ms after the corresponding trigger point (fixation point onset for PR, cue onset for EV and tone onset for CR, EV_ITI and PR_ITI). We identified the earliest pair of consecutive bins showing p < 0.01 for the respective coefficient and defined the latency as the start of the first of these bins.

Because linear models of PR_RPE and EV_RPE that included all the necessary covariates (i.e. EV, PR and CR) produced inconsistent results, we examined the encoding of these quantities using cluster analysis and population decoding as described in the text and below.

### Classification Analyses

used the logistic classifier in the Classify[] function of Mathematica with 80/20 cross validation and 100 random replications for each variable (EV, PR, CR, EV_RPE, PR_RPE). We measured accuracy as the fraction of test trials that were assigned to the correct category. We obtained the baseline level of accuracy given the label distribution for each replication by computing 10 additional classifications on the same trials but with shuffled labels. We defined the *excess accuracy* as the difference between the average classification accuracy on the real and shuffled data. Excess accuracy was estimated for each session and statistics were conducted across sessions. For neuronal responses, the input to the classification was the trial-by-trial FR of all the neurons that were simultaneously recorded in each area in that session. For licking classification analyses, the predictor was the trial-by trial LR over the time window of [−300 300] from cue onset for PR_RPE, and [0 900] from cue onset for EV_RPE. The target classes were EV_RPE or PR_RPE (signed and unsigned separately) trained using all trials.

### Noise Correlation Analyses

were computed using firing rates in the window 300ms prior to cue onset, on cells with at least 4hz mean firing rate over the duration of the trial. Since this window was before cue presentation, our measure was not contaminated by average sensitivity to EV. To account for PR sensitivity, we binned trials were binned according to PR and calculated the mean-subtracted trial by trial response to measure noise correlation.

## Acknowledgements

The work was supported by Memory and Cognitive Disorder Award from the McKnight Foundation to JG and a generous gift of surgical equipment from Synthes Inc to JG.

---

**Figure S1.**
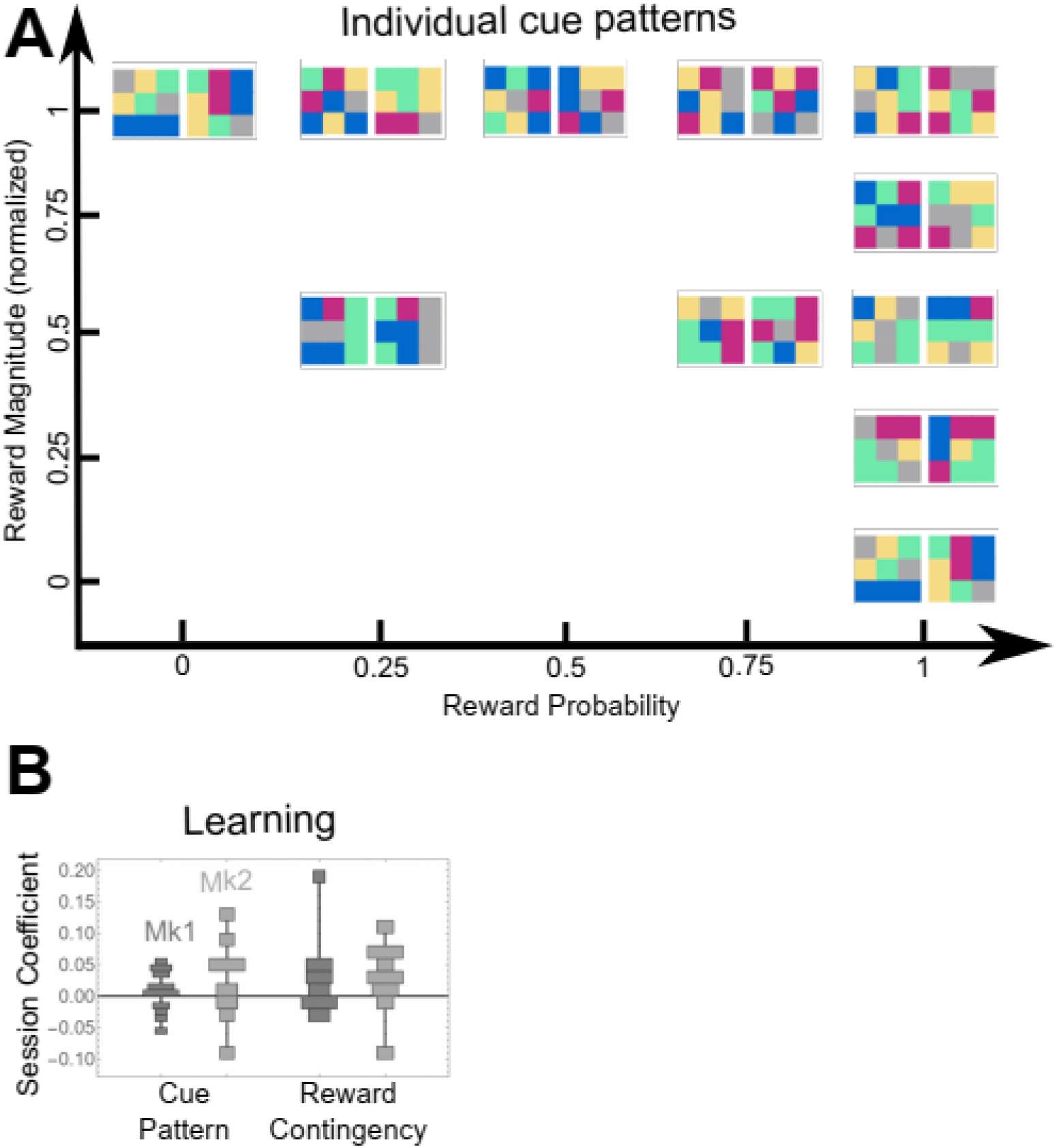
Cues and learning. (**A**) **The 20 cues and their assignment to combinations of reward magnitude and probability.** Each cue was a 3×3 colored checkerboard with each tile taking one of 5 possible colors that were defined in DKL space and isoluminant within 2cd. The distance between a pair of cues was the number of tiles that would need to be replaced for the cues to become identical. Individual cues were generated by randomly assigning colors to tiles, with the constraint that all distances are at least 4 tiles. The cues shown here were used for both monkeys. The reward mapping is shown for monkey 1 and was randomized for monkey 2 (not shown). (**B**) **Monkeys do not continuously learn about individual cues.** For each probabilistic cue (P52, P25, P57, P50, P75 in **Fig. 1B**) we computed the LR modulation based on PR (*Methods*, Eq. 1-3) restricting the trials to previous presentations of the same cue pattern or the same reward contingency. The histograms show the distribution of PR coefficients across sessions for each monkey. Had the monkeys updated the cue values the coefficients should be positive, indicating more LR after a large, versus a small, prior reward for that cue or contingencies. However, no distribution was different from 0 for the individual cues (monkey 1, p = 0.15; monkey 2, p=0.16) or reward contingency (monkey 1, p = 0.1; monkey 2, p = 0.06), showing that the monkeys were using pre-learned distributions for these highly familiar cues.

**Figure S2.**
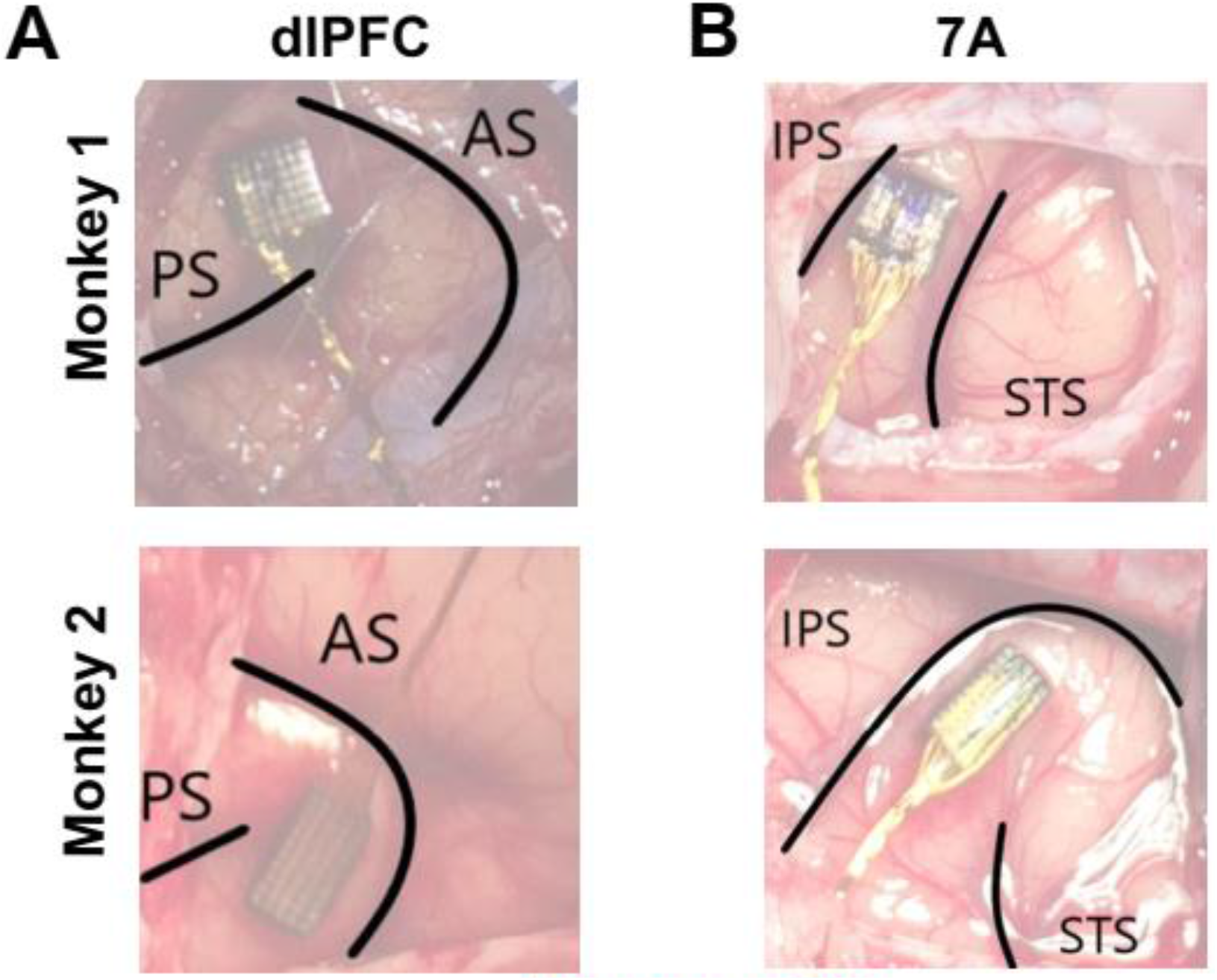
Recording sites. Intraoperative photographs showing array placements. (**A**) The dlPFC arrays were implanted between the arcuate sulcus (AS) and the principal sulcus (PS), slightly more dorsal in monkey 1 relative to monkey 2 because of vascular anatomy. (A) The 7A arrays were implanted between the intraparietal sulcus (IPS) and superior temporal sulcus (STS), in the posterior portion of this area known as OPT.

**Figure S3.**
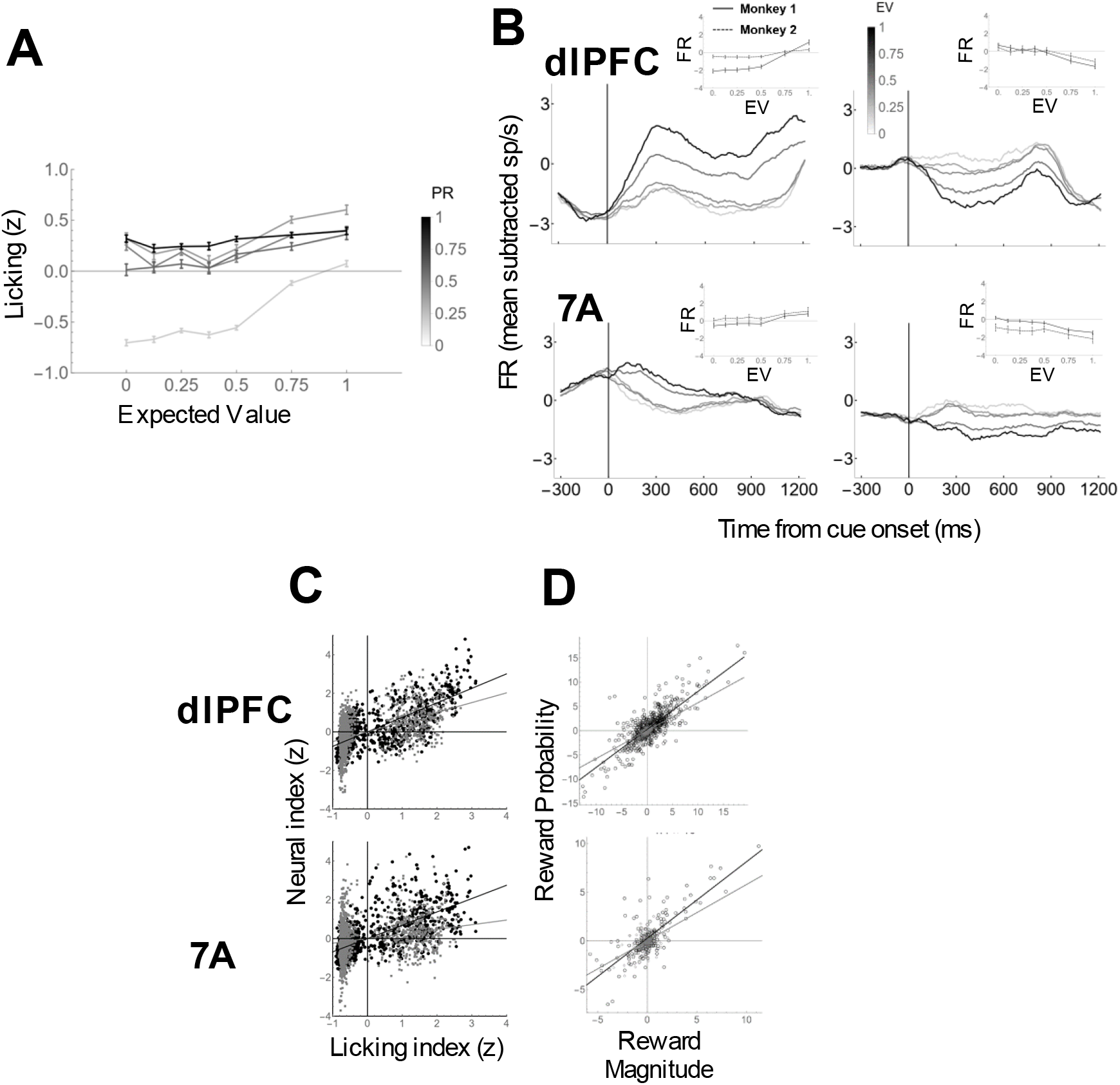
EV and PR selectivity. **(A) Anticipatory licking** (mean and SEM in the 200ms pre-tone epoch) showed strong sensitivity to PR and EV. The EV effect showed a quasi-categorical pattern, distinguishing primarily between the 2 highest and 5 lower levels of EVs. **(B) EV-sensitive cells showed categorical responses resembling the licking response.** Peri-stimulus time histograms (PSTHs; constructed by convolving raw firing rates with a 100ms boxcar moving average for 5 levels of EV) show cue-aligned firing rates for EV+ (top) and EV− (bottom) cells. The insets show firing rates (mean and SEM during the delay period) as a function of EV for each monkey. **(C) Representational similarity between licking and neural responses.** We calculated the mean licking on each trial in the 900 ms interval starting at cue onset, pooled the trials across all sessions, and partitioned the pooled dataset into 35 combinations of 7 levels of EV and 5 levels of PR. We then computed the Anderson-Darling statistic (AD) measuring the distance between the mean licking distributions in each possible pair of conditions (1,225 pairs; including those with identical contingencies). We repeated this procedure for each recorded cell using FR in the 900 ms interval starting at cue onset and pooling trials across all neurons that contributed at least 2 trials within a bin. After z-scoring each index, we calculated the correlation coefficient, across the 1,225 pairs, between the AD distances in LR and FR. Each point is a pair of contingencies and the lines show the best fit regression (black, monkey 1, gray, monkey 2. For each area and each monkey, the neural and licking representations were highly correlated (monkey 1: dlPFC r = 0.7; 7A r = 0.64; monkey 2: PFC r = 0.5; 7A r = 0.24; all p < 10^−18^). **(D) Reward magnitude and probability are highly correlated** despite occurring in separate trials, and both are highly correlated with EV (**Table 2**). Each point is one neuron and the lines show the best fit regression (black, monkey 1, gray, monkey 2)..

**FIGURE S4.**
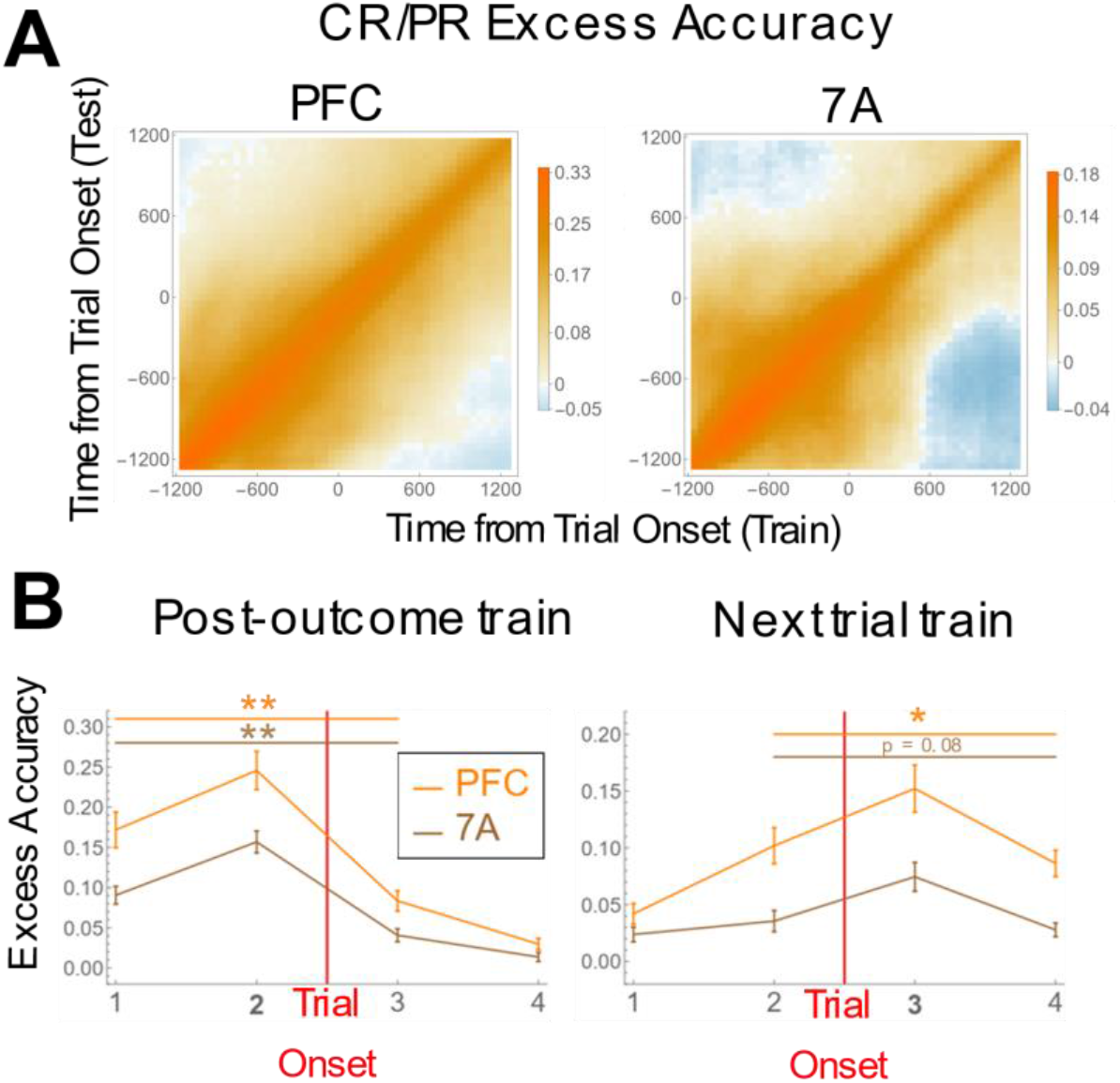
(B) PR is not a mere persistence of the CR response. **(A) Classifiers trained during the ITI do not decode reward value in the following trial and vice versa.** Classifier performance remains stable within the ITI or fixation epochs but declines at the transition between the two, suggesting that the reward representation changes at the start of a new trial. Logistic classifiers were trained on 300ms epochs, sliding at 50ms to decode the values of the current and prior rewards (CR/PR) from before to after trial onset from the network of simultaneously recorded cells in each session. These classifiers were then tested against all other epochs. The colorscale shows excess accuracy, defined as classification accuracy based on correctly labeled data minus accuracy based on data with randomly shuffled trial labels. **(B) Quantitative analysis of cross-epoch decoding.** Firing rates were divided in 4 consecutive 300 ms epochs centered on trial onset (fixation point onset). Logistic classifiers were trained to decode CR (or PR) from the activity of simultaneously recorded cells in each area using the window immediately preceding or following trial onset (left panel, window 2 vs right panel, window 3) and tested in the other time bins. The traces show classification accuracy above the shuffle control (mean and SEM across sessions) for dlPFC (orange) and 7A (brown). For windows that were equally distant from the training interval, classification was significantly more accurate when tested within the same trial relative to across trials in both areas (* p < 0.05; ** p < 0.001

**Figure S5.**
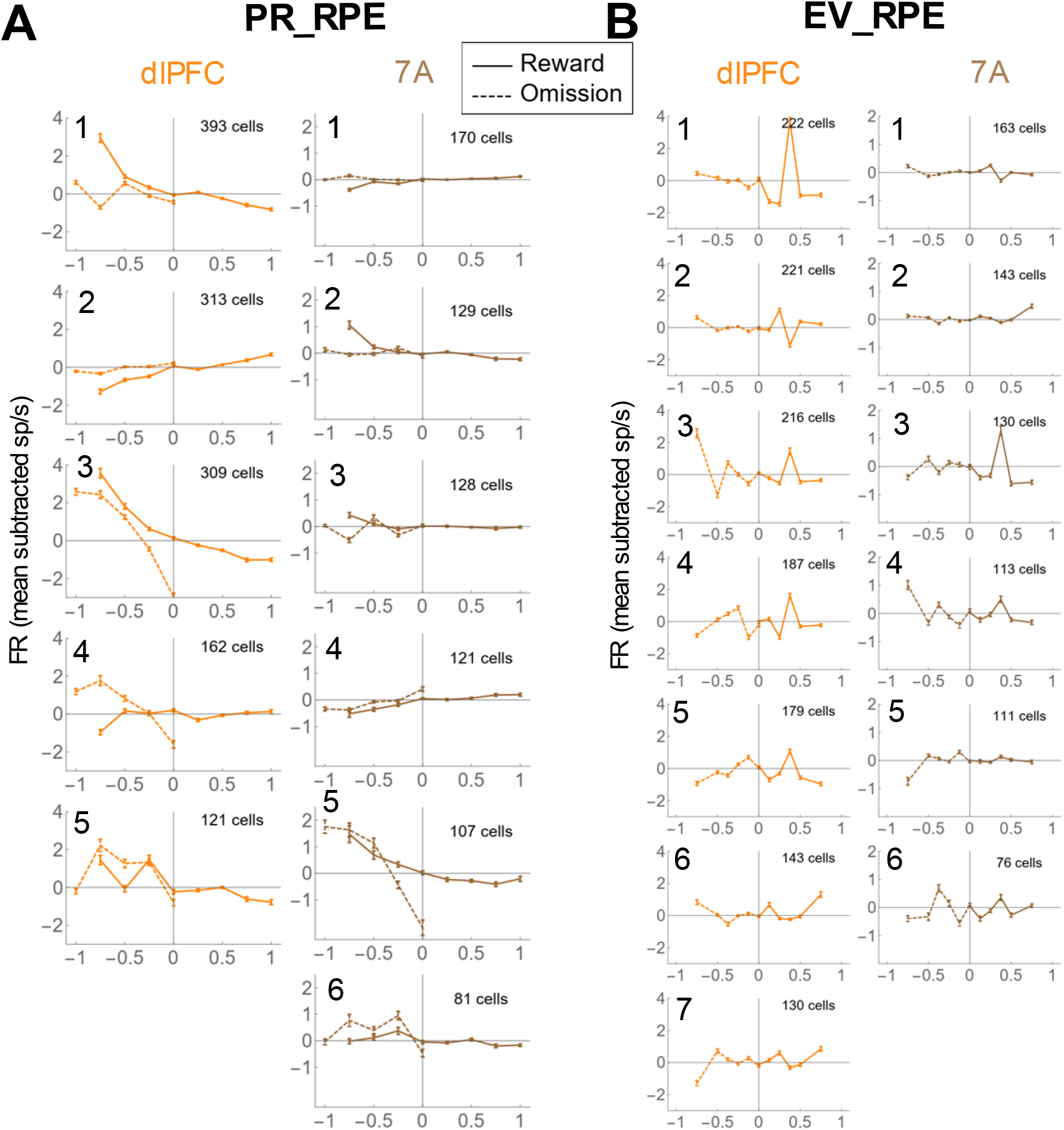
Diversity of individual neuron responses to PR_RPE (A) and EV_RPE (B) Vectors of mean deviations were constructed for each cell (as described for Fig. **6A,C**, but using the signed rather than absolute values), and then analyzed with k-means clustering with correlation-distance and k chosen for each area based on scree plots. Clusters are ordered according to size in each area. All other conventions are as in **Fig. 6A,C.**

## Notes

**Disclosure statement:** The authors declare that they have no conflict of interest.

### Competing Interest Statement

The authors have declared no competing interest.

